# Cell Morphology and Biophysical Mechanisms-Informed Traction Force Microscopy Using Machine Learning

**DOI:** 10.1101/2025.05.21.655282

**Authors:** Koki Fujiwara, Ryosuke Fujikawa, Yukihiro Suzuki, Kenta T. Suzuki, Yuichi Sakumura

## Abstract

Quantitative evaluation of cell motility is essential for understanding biological functions. Traction force microscopy (TFM) is a method for quantifying cellular traction forces. By performing inverse mathematical analysis of substrate deformations induced by cellular forces, it is possible to estimate the underlying traction forces. Among various approaches for detecting substrate deformations, the most widely used is to track the displacement of fluorescent beads randomly embedded in the substrate. However, it is well known that the accuracy of force estimation deteriorates when the observation density is low. Furthermore, standardized datasets have not been established, as validation settings—such as random force distributions on the substrate, cell size, and bead density—vary across studies. To enable quantitative evaluation, we constructed a dataset based on the biophysical mechanism by which cellular traction forces are transmitted to the substrate through stress fibers, clutch proteins, and focal adhesions. In addition, we proposed a novel machine learning model that incorporates cell shape information obtained from observations, and we designed a loss function that accounts for both force magnitude and direction. As a result, our method enables highly accurate force estimation even under sparse observation conditions.

## 1 Introduction

Cell migration and morphogenesis are fundamental to a wide range of biological processes, including neural network formation[1–4], as well as the migration of immune cells during inflammation[5, 6] and the invasion of cancer cells during tumor progression[7, 8]. These morphological transformations can be understood not only as intrinsic properties of individual cells but also as physical interactions between the cells and their surrounding environment. In particular, mechanical interactions with the substrate or neighboring cells generate forces that drive shape changes and cell locomotion[9]. Measuring such forces is essential for elucidating biological functions. To this end, various direct observation techniques have been proposed to quantify the forces that cells exert on their substrate, including atomic force microscopy (AFM)[10, 11] and micropillar arrays[12, 13]. However, AFM is limited in its field of view, making it difficult to simultaneously measure the spatial distribution of forces across the entire cell. In addition, methods based on micropillar arrays face challenges such as deformation of the substrate anchoring the pillars and interference of the structures with the cell’s natural behavior. Thus, technical limitations still remain in the direct observation of cellular forces.

TFM[14] is a widely used method to estimate the forces that cells exert on their substrate. TFM tracks substrate deformations induced by cells on a compliant substrate and inversely estimates the corresponding force field. Among the various techniques for detecting such deformations, the most common involves tracking the displacement of fluorescent beads randomly embedded in the substrate [15–18][Fig. 1 A]. This method is compatible with conventional optical microscopy and is technically straight-forward to implement. However, in bead-based TFM, the spatial resolution of the displacement field strongly depends on the bead density, which directly affects the accuracy of force estimation. When the bead density is low, insufficient displacement information leads to a significant reduction in estimation accuracy[19]. Conversely, increasing the bead density can alter the mechanical properties of the substrate and influence cellular behavior, making it difficult to achieve in practice[20, 21]. To address these challenges, Bayesian inference methods[18, 22, 23] and machine learning approaches[24–26] have been explored as alternatives to maintain high accuracy under low-density conditions.

**Figure 1.**
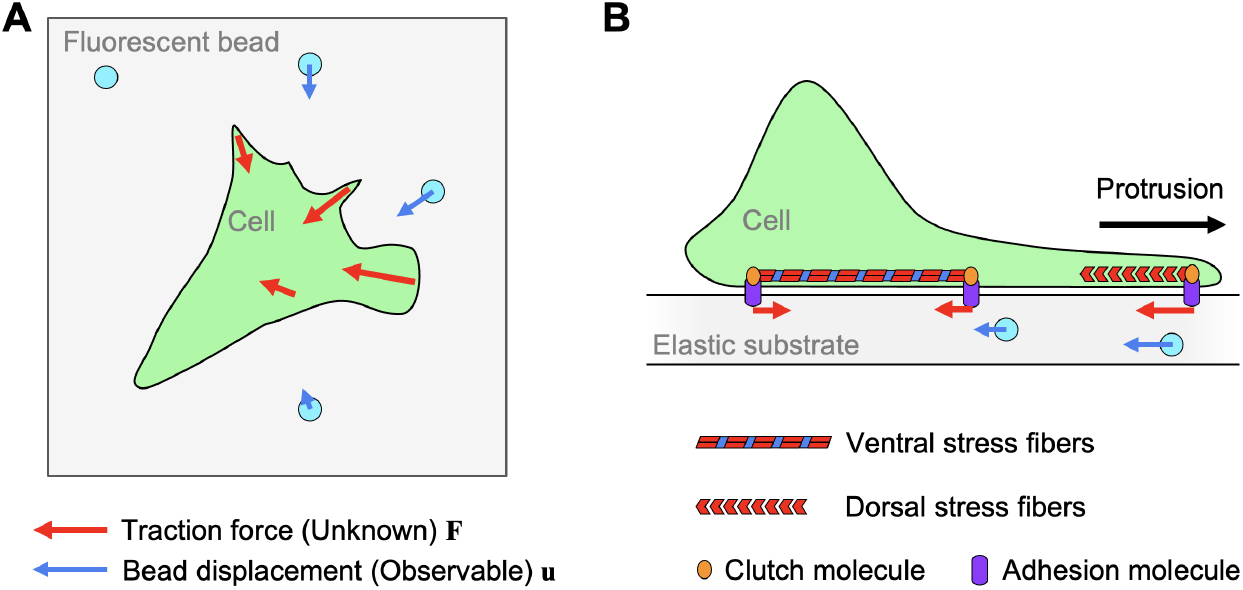
Diagram of Cellular Traction Force Estimation. (A) Overview of traction force microscopy (TFM). This illustrates how a cell on a substrate (green) exerts forces (**F**; red arrows) that cause the displacement of fluorescent beads (**u**; blue circles and arrows) embedded on the substrate surface. By inversely analyzing these bead displacements, the traction forces can be estimated. (B) Cellular adhesion is known to be mediated by two types of stress fibers: ventral stress fibers and dorsal stress fibers, which are connected to the substrate via clutch molecules. Contraction and polymerization of these fibers generate inward forces toward the cell center. As a result, the cell maintains adhesion to the substrate while migrating forward (black arrow).

We previously proposed a Bayesian force estimation method [23] incorporating prior distributions based on biological insights that cellular forces tend to be directed inward near the cell edge[16, 27, 28]. By integrating this prior knowledge with an appropriate inference algorithm, we achieved improved accuracy in low bead-density settings. Nevertheless, the displacement data of fluorescent beads is often contaminated by observational noise due to limitations in microscope resolution and tracking algorithms. Moreover, the accuracy of the boundary-dependent force prior can deteriorate in the presence of complex or noisy cell shape data. As a result, our previous method still faced challenges in terms of robustness against bead displacement noise and cell shape irregularities.

Recently, the application of machine learning in biological research has rapidly progressed, and TFM is no exception. In particular, convolutional neural networks (CNNs), which excel at extracting local features and preserving spatial information, have been introduced for traction force estimation. These data-driven approaches learn the relationship between substrate deformation and bead displacement, enabling inverse estimation without reliance on physical models. As a result, they offer advantages such as computational speed, flexibility, and robustness to noise. Once trained, CNN models can instantly output force distributions from input data, offering practical utility.

However, existing machine learning approaches have primarily been evaluated under ideal conditions involving large cells and high bead densities. Their performance under more challenging scenarios—such as small cells or low-density observations—remains insufficiently verified. Furthermore, most CNN-based methods have yet to incorporate biological insights such as the relationship between cell morphology and force distribution.

In this study, we investigate the effect of incorporating cell shape information into the force estimation model, focusing on biologically plausible mechanisms underlying force generation. To solve the inverse elasticity problem, we employ a CNN-based method with high predictive reliability. Specifically, we provide the CNN with three input channels: the x-component of bead displacement, the y-component of bead displacement, and a binary mask indicating the interior and exterior of the cell. A U-Net architecture[29] is used for the learning process. Large-scale synthetic training data are generated by applying a generative model to cell images obtained from public databases, yielding binary masks of cell shapes. Within each generated cell shape, realistic force distributions are assigned based on the magnitude of traction forces and directions obtained using level-set methods (LSM) and mean curvature flow (MCF)[30–32], together with additional rules reflecting the physical mechanisms of force generation compatible with the geometry of the cell. The resulting bead displacements are then computed by numerically solving the elastic equation for various bead densities and random bead placements. Using these synthetic datasets, the model is trained to minimize the error between the predicted and ground-truth traction forces, based on both displacement and shape inputs.

To evaluate the contribution of shape information, we compare models trained and tested with and without the cell shape channel, using the same CNN architecture. Our results demonstrate that incorporating shape information improves the accuracy of traction force estimation compared to models that rely solely on bead displacements. Notably, under low bead-density conditions, shape-aware models maintain stable predictions and prevent the severe accuracy degradation observed in shape-agnostic models, demonstrating robust performance against both displacement noise and cell shape uncertainties. Additionally, further improvements are anticipated through refined model designs tailored to the spatial characteristics of traction forces. This study demonstrates that accurate force estimation is achievable even under low-resolution or low-density observation conditions, where conventional TFM approaches tend to fail, and is expected to provide a foundation for future non-invasive biomechanical analyses.

## 2 Methods

### 2.1 Cell Shape Data Acquisition

To construct a machine learning-based traction force estimation model, an appropriate training dataset is essential. The input to the model consists of three channels: the x- and y-components of the bead displacement, and a binary mask indicating the interior and exterior of the cell. Since the spatial distribution of cellular traction forces is known to depend on the cell’s morphology and contact mode with the substrate, realistic cell shape information is necessary not only to determine the force magnitude and direction, but also to define the spatial origin of force generation.

In this study, we used the publicly available Oriented Cell Dataset (OCD) published on IEEE DataPort[33], which includes microscopy images obtained from real observations. This dataset contains five types of cell lines: A172 and U251 (human glioblastoma), MCF7 (human breast cancer), MRC5 (human lung fibroblast), and SCC25 (human squamous cell carcinoma).

Each image in the dataset contains multiple cells, which are not directly suitable for machine learning input. Therefore, we applied Cellpose[34], a specialized segmentation tool for cell images, to separate individual cells. Based on the segments extracted by Cellpose, we constructed binary masks that clearly distinguish between the inside and outside of each cell. These masks were then used as input for downstream processing and as part of the training data for the machine learning model.

### 2.2 Cell Shape Generation Using Generative Models

Accurate traction force estimation via machine learning requires a training dataset that adequately reflects the diversity of cell shapes. However, the morphological variations captured through real microscopy observations are limited, and insufficient to ensure both the quantity and diversity necessary for robust model training.

To address this, we applied data augmentation using a generative approach. Specifically, we used a Deep Convolutional Generative Adversarial Network (DCGAN)[35] to generate new cell shapes that statistically reflect the morphological distribution of MRC5 cells, one of the cell types included in the OCD.

DCGAN offers the advantage of being trainable in an unsupervised manner without the need for label information, allowing efficient use of data. Compared to recently popular diffusion-based generative models, DCGAN is capable of producing high-quality binary masks with greater computational efficiency and faster inference speed. For this reason, it was deemed suitable for the present study, where the goal is to generate binary cell masks.

Using the trained generative model, we synthesized a large number of artificial cell shapes. These shapes were formatted into binary masks and incorporated into the training dataset. As a result, we constructed a dataset that maintains the biological reliability of real images while significantly expanding the morphological diversity.

### 2.3 Traction Force Modeling Framework

In constructing a traction force estimation model, it is essential to explicitly define the relationship between cell morphology and the expected traction force field during training. In our previous work, we adopted an approach that incorporates biologically informed priors for the magnitude and direction of traction forces. Specifically, we utilized the fact that actin polymerization is active near the cell boundary, resulting in large inward-directed traction forces. Based on this prior knowledge, we modeled the traction force vector field using models for the magnitude and LSF with MCF.

However, beyond the magnitude and direction, the location where traction forces are generated is also a critical component. Despite this, conventional modeling approaches have often neglected this spatial aspect. This is a significant limitation, especially in light of biological evidence showing that the spatial distribution of traction forces is influenced by the interaction between cell shape and its contact with the substrate.

To address this issue, we propose a new modeling approach that incorporates not only the magnitude and direction but also the biologically plausible locations of force generation. This results in a more comprehensive force distribution model that maintains both biological consistency and mathematical coherence. Below, we describe each component in detail.

#### Step 1: Modeling the Force Magnitude

The magnitude of traction force is known to decay with distance from the cell boundary. Specifically, force tends to be strongest near the boundary, where actin polymerization is active, and gradually decreases toward the interior. Following this observation, we modeled the magnitude *m*(**x**) as a Gaussian-like function of the distance *r*(**x**) from the cell boundary:

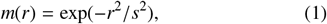

where *s*^2^ is a hyperparameter set in proportion to 10% of the cell area, enabling scale adjustment depending on the cell size. This formulation captures the spatially localized nature of traction forces with a maximum at the boundary and exponential decay toward the interior.

#### Step 2: Modeling the Force Direction

The direction of traction forces is determined based on the biological observation that actin polymerization drives inward force near the cell edge. We define the direction vector field **d**(**x**) using the normalized gradient of the LSF *ϕ*(**x**, *t*) that represents the cell shape:

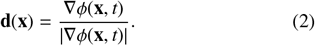

The initial shape *ϕ*(**x**, *t* = 0) contains protrusions and regions of high curvature along the cell boundary, which often cause the gradient vector field to become discontinuous and unstable. To address this issue, we employed MCF to smoothly deform the shape and used the edge at time *t* = *τ*. In this study, *τ* was set such that *τ* > 0.1 [cell area]/*D*, based on 10% of the cell area, where *D* is the diffusion coefficient associated with MCF. This approach enables the construction of a traction force direction vector field that preserves the complexity of the cell boundary while achieving both biological fidelity and mathematical smoothness.

#### Step 3: Modeling the Force Generation Locations

Although not all aspects of cellular force transmission are fully understood, it is well known that migrating cells form pseudopodia at adhesion sites with the substrate and exert localized forces at these sites[36]. These forces are transmitted through structures known as focal adhesions, which serve as mechanical transmission points formed between the cell membrane and the substrate.

Focal adhesions are protein complexes that connect actin filaments to the substrate and are composed of various proteins, including integrin, talin, and vinculin, which are sequentially involved in their assembly[37]. These structures are functionally associated with actin stress fibers, of which three major types exist in cells: ventral stress fibers, dorsal stress fibers, and transverse arcs[38].

Ventral stress fibers are contractile actin bundles that are anchored at both ends to focal adhesions. They span the basal surface of the cell and play a central role in transmitting traction forces to the substrate through actomyosin-driven contraction. In contrast, dorsal stress fibers are anchored at only one end to a focal adhesion, typically located near the cell periphery, while their other end extends into the cytoplasm. These fibers are primarily involved in cell elongation and pseudopodia formation, and serve as structural guides for organizing other cytoskeletal components. Transverse arcs, on the other hand, are curved actin bundles located within the central region of the cytoplasm. Unlike the other two types, they are not directly connected to focal adhesions, but instead contribute to the mechanical stability of the cytoskeleton by dynamically interacting with both ventral and dorsal fibers.

Thus, the formation of pseudopodia and focal adhesions, both essential for traction force generation, is functionally coordinated with ventral and dorsal stress fibers. Based on this structural and functional coordination, we model the locations at which traction forces are generated.

For ventral stress fibers, we used the LSF, denoted as *ϕ*(**x**, *t*), and specifically employed its state at time *t* = *τ*, i.e., *ϕ*(**x**, *t* = *τ*). Based on this, representative points were randomly selected from the region where *ϕ*(**x**, *t* = *τ*) < 0, representing substrate-contacting regions prior to morphological change. To reproduce adhesion patches, along with the 8-neighboring pixels around each selected point. To avoid the influence of cell size, the total patch area was adjusted to be approximately 10% of the intracellular region defined by *ϕ*(**x**, *t* = *τ*).

Dorsal stress fibers are primarily distributed near the peripheral region of the cell. First, based on the LSF *ϕ*(**x**, *t* = 0), we extracted contour points with high curvature variation, which correspond to protrusive structures such as pseudopodia and filopodia. Next, we set the time *τ*^′^ such that *τ*^′^ > 0.01 [cell area]/*D*, and focused on the region enclosed by *ϕ*(**x**, *t* = 0) and *ϕ*(**x**, *t* = *τ*^′^). Within this region, we selected adhesion sites for dorsal stress fibers only if the nearest point on *ϕ*(**x**, *t* = 0) belonged to a high-curvature contour point.

By incorporating both partial structural dependence and randomness into the overall cell morphology and local structural characteristics, we modeled the biological diversity of adhesion site distribution.

### 2.4 Computation of Bead Displacement

When a cell exerts traction force **f**(**x**) on the substrate, fluorescent beads embedded on the substrate surface undergo a displacement **u**(**x**). This displacement is described as a convolution integral based on the Boussinesq Green’s function 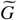 [39]:

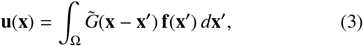

where Ω denotes the defined domain on the substrate, and 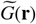 is the Green’s function for a semi-infinite elastic medium subjected to surface traction,

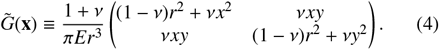

Here, the vector **r** = **x** − **x**′, and *r* = |**r**| is the Euclidean norm. The Green’s function is parameterized by the elastic modulus Young’s modulus *E* and the Poisson’s ratio ν of the substrate.

Since solving the continuous convolution integral in Eq.As shown in [Eq. 3] is computationally intractable, we discretize the domain *U* into *N* square lattice elements. We assume a constant traction force **f**(**x**′_*n*_) within each element, leading to the following discretized approximation:

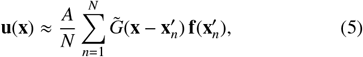

where *A* represents the total area of the cellular region, so *A*/*N* corresponds to the area per discrete element. This formulation results in a linear system that connects the traction force field to bead displacement through a physically interpretable model grounded in elasticity theory.

### 2.5 Incorporation of Noise in Bead Displacement

In practical TFM experiments, measurements of bead displacement are inevitably subject to observational noise arising from several factors such as limited spatial resolution of microscopy, fluctuations in fluorescence intensity, and errors in particle tracking during image processing. These uncertainties are reflected in the displacement vector **u**(**x**) as additive noise.

To emulate these experimental uncertainties and promote robustness of the learning model to measurement noise, we add Gaussian-distributed noise to the synthetic bead displacement data. The perturbed displacement is modeled as:

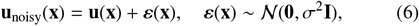

where the standard deviation of the noise, σ, is set to be approximately equal to the resolution of the spatial grid (grid interval). This ensures that the noise scale naturally follows the scaling of the coordinate system, thereby maintaining consistent noise characteristics across conditions with different cell sizes or image scales.

### 2.6 Machine Learning Model and Loss Function

For traction force estimation, we adopted the U-Net architecture, which is widely used in image processing tasks. U-Net captures both local and global spatial features from input images and produces spatially resolved outputs, making it well suited for the inverse mapping in TFM. Its effectiveness has been previously demonstrated in TFM applications, and we followed this precedent in our model design.

The input to our model consists of three channels: the x- and y-components of the bead displacement vector, along with a binary mask indicating the cell interior and exterior. The output comprises two channels corresponding to the x- and y-components of the traction force vector. To evaluate the impact of morphological information on the accuracy of traction force estimation, we also constructed a comparative model that uses the same architecture but omits the binary mask. This baseline model accepts only the x- and y-components of bead displacement as input, while maintaining the same two-channel output format. This design allows us to isolate and quantify the influence of cell shape information on model performance.

While we made no major changes to the U-Net architecture itself, we designed a specialized loss function that reflects the vectorial nature of traction forces. In general, machine learning models commonly employ the Mean Squared Error (MSE) to minimize the discrepancy between predictions and ground truth. However, MSE primarily evaluates the difference in vector magnitudes, which is insufficient when dealing with traction forces that have both magnitude and direction.

To address this, we introduced an additional loss term based on cosine similarity between the predicted and true traction vectors[40]. The combined loss function is defined as follows:

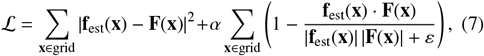

where **f**_est_(**x**) denotes the estimated traction force vector, and **F**(**x**) is the corresponding ground truth vector. The first term minimizes the squared norm difference, representing the error in force magnitude. The second term penalizes directional discrepancies between the predicted and true vectors, with *α* controlling the weight of this directional term. A small constant *ε* is added to the denominator to avoid division by zero.

By combining magnitude- and direction-based error measures, this loss function enables more physically meaningful learning that reflects the true nature of traction forces. Consequently, it facilitates the development of more accurate estimation models.

### 2.7 Model Training and Evaluation Protocol

To comprehensively evaluate the performance of the proposed traction force estimation model, we conducted experimental validations from three key perspectives:

**Figure 2.**
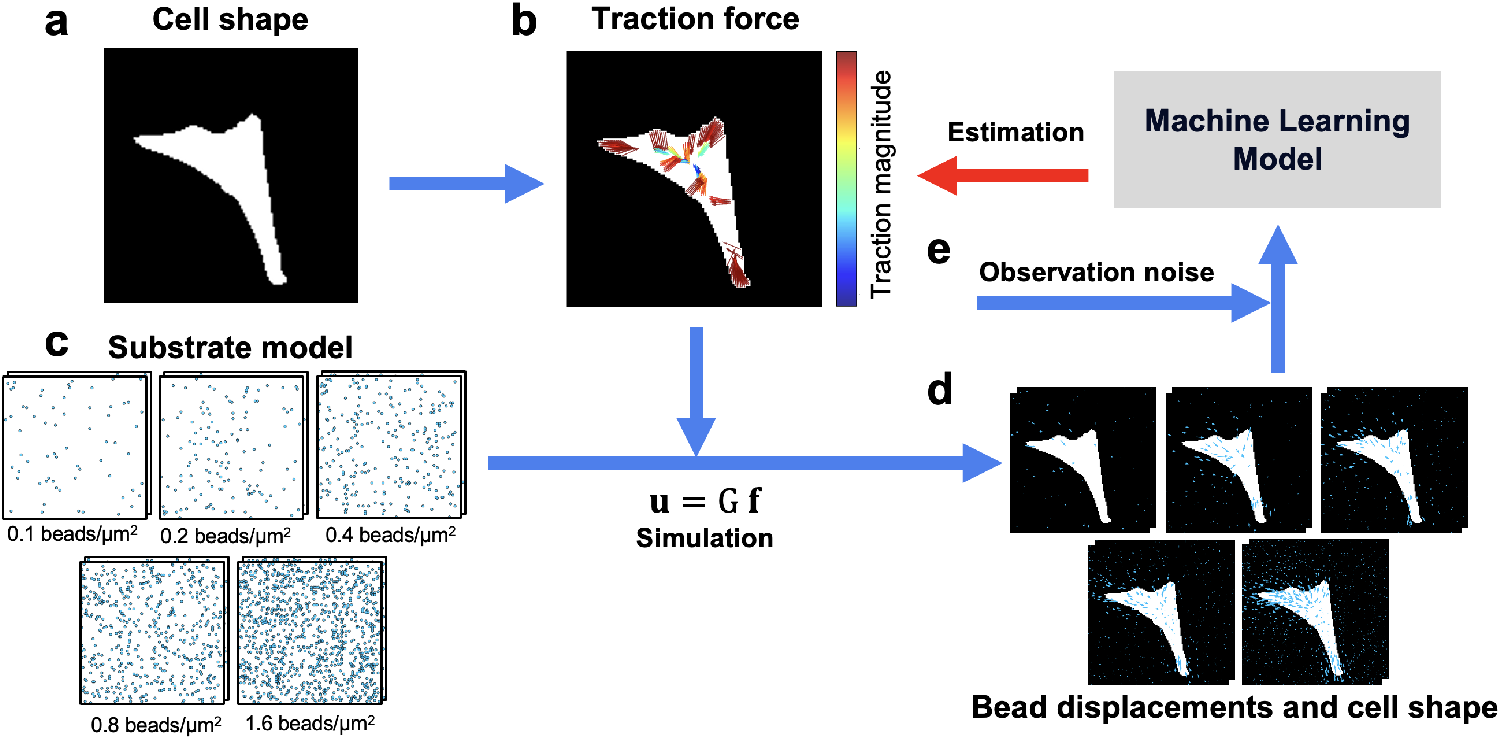
Workflow for estimating traction forces from synthetic datasets. (a) Cell shape masks were prepared, and (b) biologically inspired traction force distributions were generated within the cell area. The direction of the forces is indicated by the orientation of the arrows, and the magnitude is represented by both the length and color of the arrows. (c) Five different bead densities and two types of random bead distributions were placed on the substrate. (d) Bead displacements were computed using the Boussinesq approximation, **u** = **Gf**. (e) Observation noise was added to the displacement data using a 2D Gaussian distribution with a standard deviation of 0.2 *μ*m. Finally, the noisy displacement data and the cell shape information were input into a machine learning model, which was trained to estimate the underlying traction force distribution.

#### Generalization Performance Using Synthetic Datasets

To assess the influence of bead density on estimation accuracy under controlled conditions, we designed experiments using fully synthetic data. The training dataset was constructed using synthetic traction force fields and bead displacement fields generated based on real cell shape masks. All inputs and outputs were resampled to 128 × 128 grids to conform with the U-Net architecture. The grid resolution was set to match the spatial resolution of a confocal laser scanning microscope (0.2 μm per pixel), enabling consistent scaling across different cell sizes and bead densities.

For training, we used 4,800 cell shape masks generated by a DCGAN trained on MRC5 cells. For each mask, we simulated bead displacement data under five bead density conditions: 0.1, 0.2, 0.4, 0.8, and 1.6 beads/μm^2^. Additionally, two random bead placements were generated for each density condition, resulting in a total of 48,000 synthetic data samples (4,800 × 5 × 2).

For testing, we selected 100 cell shapes randomly from previously unseen cell types (A172, U251, MCF7, SCC25). For each shape, traction forces and bead displacements were generated under the same five bead density conditions, resulting in 500 test samples. Model training was conducted using a learning rate of 0.0001, a batch size of 128, and a maximum of 100 epochs. The model corresponding to the epoch with the lowest validation error was selected for evaluation.

#### Impact of Morphological Information on Model Performance

To evaluate the effect of incorporating cell shape information into the model input, we compared two configurations using the same U-Net architecture: with cell shape mask (three-channel input: bead displacement in x and y directions + shape mask) and without shape mask (two-channel input: bead displacement only). Both models were trained and evaluated independently. This design allowed us to rigorously assess the contribution of cell shape information to the spatial accuracy of traction force estimation.

#### Influence of Loss Function Parameters

Finally, we investigated the impact of incorporating directional information into the loss function. The proposed loss function is a linear combination of the MSE for force magnitude and a cosine similarity-based loss for force direction. We adjusted the weight parameter *α* to control the contribution of angular information, and trained models under four different conditions: *α* = 0 (MSE only), 0.05, 0.1 (equal weighting), and 0.5.

Through this comparison, we quantitatively evaluated how directional information affects the spatial accuracy of the estimated traction field. By structuring our evaluation across three axes—dataset configuration, input information, and loss function design—we clarified the influence of each modeling choice on traction force inference performance.

### 2.8 Evaluation Metrics

To quantitatively assess the performance of the traction force estimation model, we employed seven evaluation metrics covering three aspects of traction force fields: magnitude, direction, and spatial distribution. These metrics were designed to capture both local errors at biologically meaningful regions (e.g., adhesion and background zones) and global estimation accuracy of the entire force vector field. By leveraging cell shape masks to distinguish between intracellular and extracellular background regions, we were able to obtain more spatially localized performance measures than conventional approaches.

**Figure 3.**
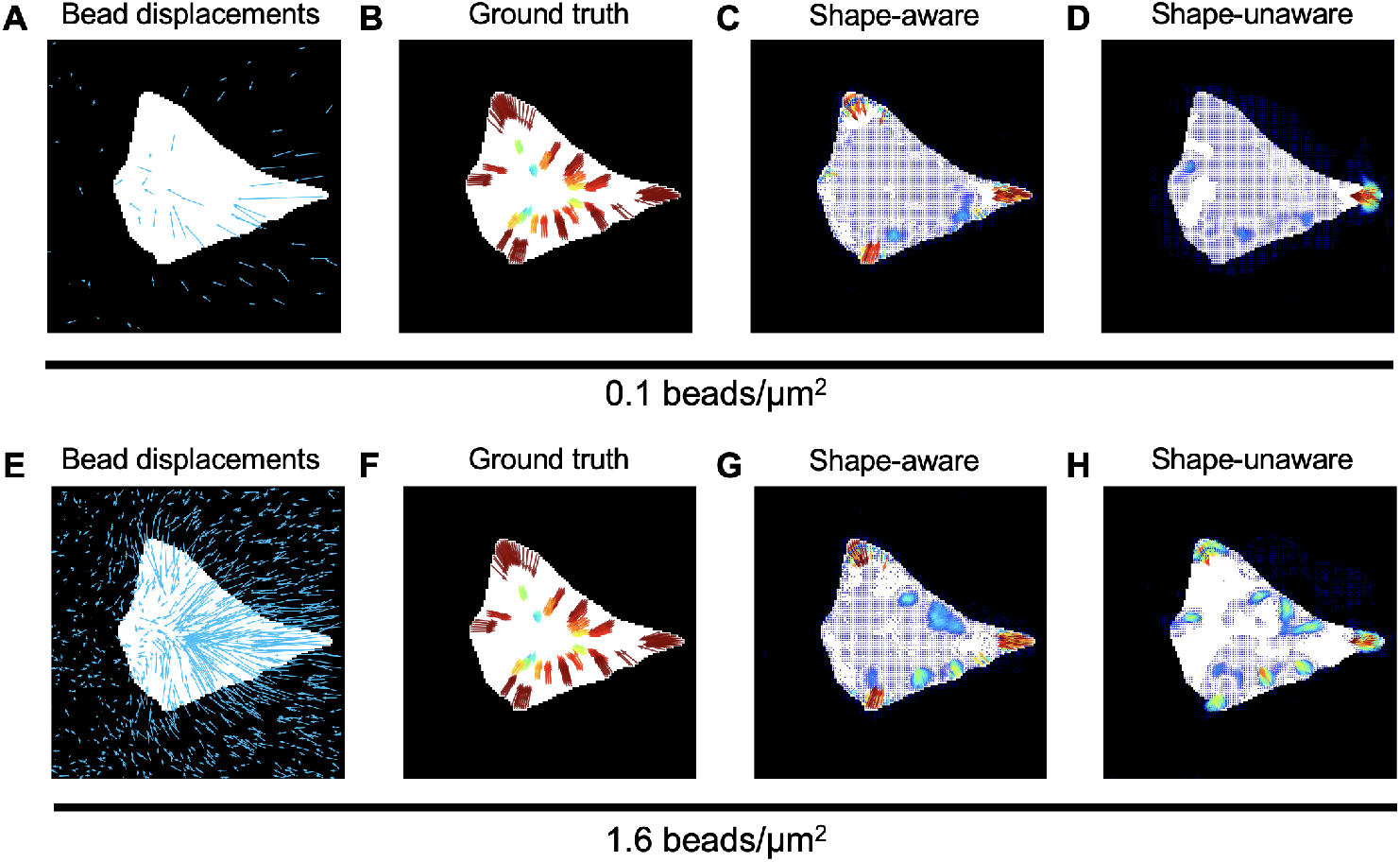
Comparison of traction force estimation results under different bead densities and trained models. (A) Blue lines represent the displacements of randomly distributed beads on the substrate model (density: 0.1 beads/*μ*m^2^). (B) Synthetic traction forces. The direction of the forces is indicated by the orientation of the arrows, and the magnitude is represented by both the length and color of the arrows. (C) Estimated results using a model trained with cell shape information and using only the MSE loss function. Cell shape information was also provided during inference. (D) Estimated results using a model trained without cell shape information but still using only the MSE loss function. Cell shape information was still provided during inference. (E) Blue lines represent the displacements of randomly distributed beads on the substrate model (density: 1.6 beads/*μ*m^2^). (F) Synthetic traction forces, identical to (B), used to compare the effect of bead density. (G, H) Same estimation as (C, D), but using a bead density of 1.6 beads/*μ*m^2^ and models trained using only the MSE loss function. In all calculations, the estimated force distributions were resampled on a 128 × 128 grid. For clarity, force vectors with magnitudes less than or equal to 0.01 were omitted in (C), (D), (G), and (H).

**f**_est_(**x**) denotes the estimated traction force vector, and **F**(**x**) represents the corresponding ground truth traction force vector. 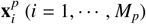 indicates the positions where traction forces are actually present, while 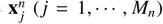 represents the positions where no traction forces occur. Furthermore, 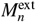 and 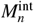 denote the number of locations without traction force in the extracellular and intracellular regions, respectively.

*Deviation of Traction Direction at Adhesions (DDA)*[26]

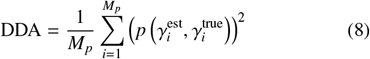

*Deviation of Traction Magnitude at Adhesions (DTMA)*[22]

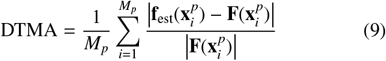

*Absolute Deviation of Traction Magnitude at Adhesions (ADTMA)*[26]

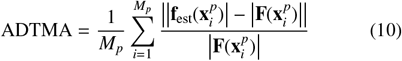

*Mean Squared Error (MSE)*

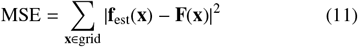

*Deviation of Traction Magnitude in the Background (DTMB)*[22]

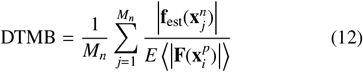

*Deviation of Traction Magnitude in the Extracellular Background (DTMEB)*

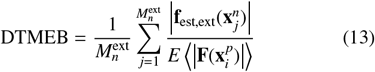

*Deviation of Traction Magnitude in the Intracellular Background (DTMIB)*

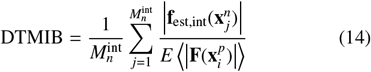

## 3 Results

### 3.1 Robustness of Cell Shape-Aware Models Under Low Bead Density

We evaluated traction force estimation performance using synthetic datasets with varying bead densities, comparing two models: one with cell shape information included as input (shape-aware model) and one without (shape-unaware model). Both models were tested on identical cell shapes and traction patterns under five bead density conditions ranging from 0.1 to 1.6 beads/μm^2^.

Quantitative comparisons were conducted using multiple evaluation metrics targeting force magnitude, direction, and spatial distribution. As shown by DTMA and ADTMA, the shape-aware model consistently outperformed the shape-unaware model in low-density conditions (0.1–0.4 beads/μm^2^), while the latter showed a notable drop in accuracy. These results indicate that incorporating morphological information enables robust generalization to previously unseen cell shapes.

The same trend was observed with MSE, though the interpretation of MSE differs. Because MSE aggregates errors across the entire image regardless of force locality, it can sometimes be deceptively low when the model outputs widespread weak forces. In contrast, DTMA and ADTMA better capture localized deviations at force-generating sites. These findings highlight the effectiveness of shape-aware modeling particularly in low-information regimes.

### 3.2 Suppression of Erroneous Predictions in Non-Adherent Regions

In addition to reproducing traction force distributions in adherent regions, it is equally important for models to suppress predictions in background regions where no force is expected. Biologically, traction forces should only occur where the cell is physically attached to the substrate. Therefore, false predictions in non-adherent regions are biologically implausible and should be evaluated separately.

To quantify such errors, we first used DTMB, a conventional metric for evaluating background noise. Furthermore, by leveraging cell shape masks, we subdivided the background into extracellular and intracellular zones and introduced two new metrics: DTMEB and DTMIB.

DTMB results showed little difference between models at high bead densities (0.8, 1.6 beads/μm^2^). However, at lower densities (0.1, 0.2 beads/μm^2^), the shape-aware model demonstrated slightly better suppression of background noise. In contrast, DTMIB revealed that the shape-unaware model slightly outperformed the shape-aware model in suppressing noise within the intracellular region, possibly due to its lower sensitivity to boundary-localized signals.

DTMEB analysis showed a clear advantage of the shape-aware model in reducing erroneous force predictions in extra-cellular regions. The shape-unaware model exhibited increased background noise as bead density decreased. The introduction of DTMEB and DTMIB allowed for finer-grained localization of background error sources, which were indistinguishable in conventional DTMB evaluations. Overall, these results suggest that the inclusion of shape information contributes to improved biological plausibility by minimizing false-positive outputs outside the cell.

### 3.3 Effect of Cosine Similarity in the Loss Function

To improve directional accuracy, we introduced a cosine similarity term into the loss function, alongside the standard MSE for magnitude error. Four different weight settings for the cosine term were tested: *α* = 0 (MSE only), 0.05, 0.1 (equal weighting), and 0.5.

Analysis of DDA, a metric for directional error, showed that increasing *α* consistently led to lower DDA values, indicating improved alignment of predicted and true traction vectors. This trend was observed regardless of bead density.

However, incorporating cosine similarity was not universally beneficial across all metrics. For DTMA and ADTMA, which focus on force magnitude, the effect of varying *α* was minimal. Interestingly, metrics related to background suppression (DTMB and DTMEB) were sensitive to loss weighting. The shape-aware model achieved the best performance under balanced loss weighting (*α* = 0.1), whereas the shape-unaware model showed no clear improvement. When *α* = 0.5, we observed that directional accuracy improved, but this came at the cost of increased low-magnitude noise in non-force regions.

These findings suggest that including directional terms in the loss function enhances angular accuracy but may introduce false predictions in background areas, highlighting a trade-off between directional precision and biological plausibility.

## 4 Discussion

In this study, we proposed a novel machine learning framework that integrates cell morphology information to enhance the accuracy of traction force estimation. Since cell shape can be readily observed in the TFM process without additional manipulation, it offers a practical and biologically meaningful input. Importantly, cell morphology reflects the mechanical history and spatial distribution of intracellular forces due to both endogenous activity and external stimuli. Incorporating such morphological information provides a new physical prior, complementing bead displacement data and increasing the information content available to the model. Comparative analysis revealed that models incorporating cell shape information achieved higher estimation accuracy than those relying solely on bead displacement.

Achieving high-accuracy inference through machine learning requires not only suitable model architectures, but also carefully designed training datasets. When datasets lack sufficient variability in cell morphology or bead density, the model may fail to generalize to unseen shapes or information-sparse conditions. To overcome this limitation, we employed a DCGAN to generate a wide range of synthetic cell shapes and systematically varied bead densities during training. As a result, the model exhibited robust generalization performance even under low-density and shape-unseen scenarios, suggesting that including diverse and realistic training conditions is essential for achieving strong generalization.

Moreover, to fully exploit the potential of the learning model, the design of the loss function must align with the physical nature of the target. Traction force is a vector quantity, and minimizing magnitude error alone using MSE is insufficient. We therefore introduced an additional loss term based on cosine similarity to capture directional accuracy. This enhancement improved both angular precision and background suppression. However, we also observed a trade-off: overemphasizing angular accuracy led to increased spurious outputs in non-force regions. These findings suggest that loss function design should be tailored according to the target application, balancing priorities between magnitude, direction, and noise suppression.

**Figure 4.**
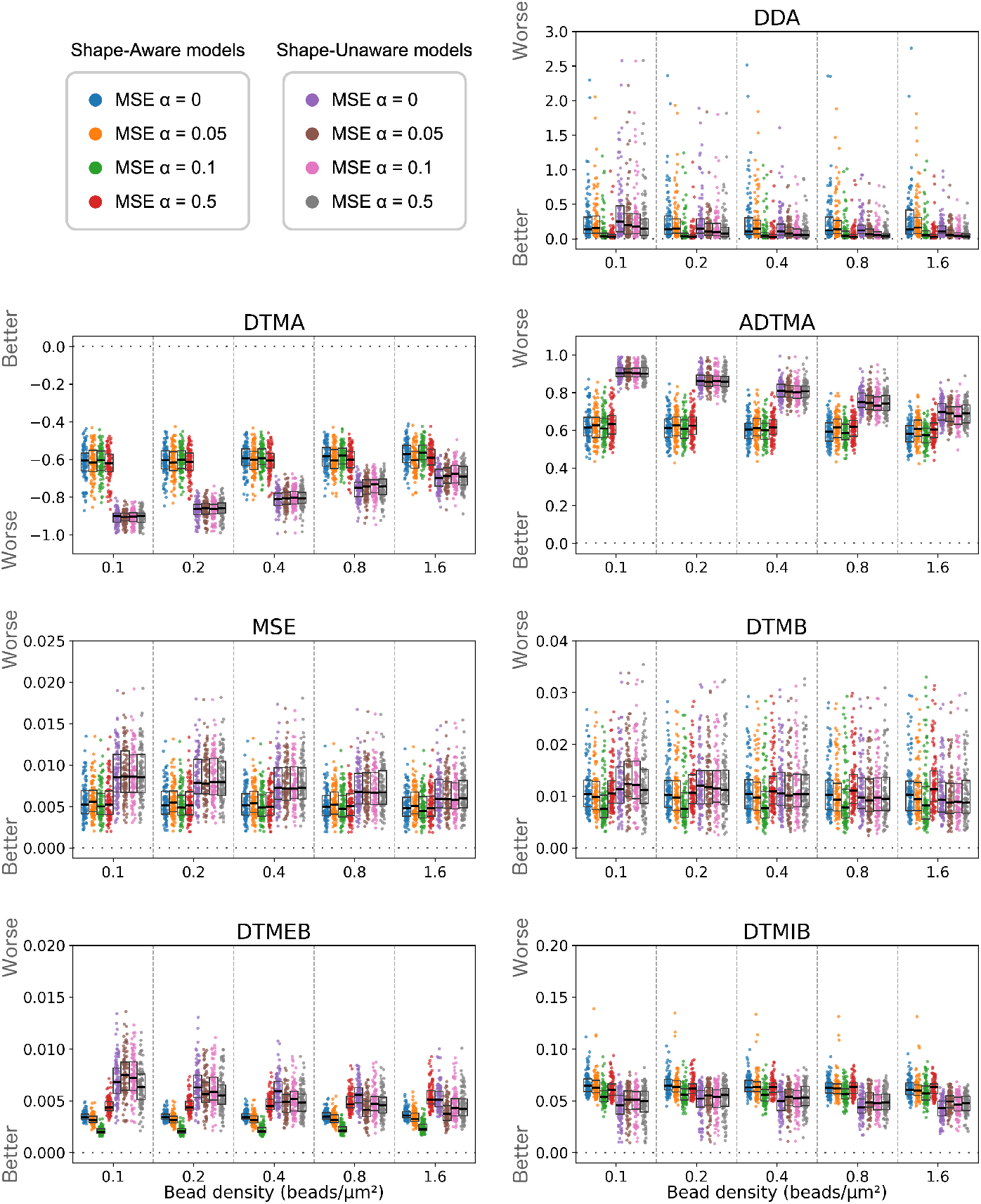
Comparison of seven evaluation metrics across different bead densities, model types, and loss function settings. Each plot shows the performance of models trained under eight conditions: two model types (shape-aware and shape-unaware) and four values of *α* for the loss function (0, 0.05, 0.1, and 0.5). DDA: Represents the deviation of traction force direction at adhesion sites. The value is expressed in radians. DTMA: Represents the deviation of traction force magnitude. ADTMA: Represents the absolute deviation of traction force magnitude. MSE: Represents the mean squared error across the entire prediction grid. DTMB: Represents the deviation of traction force magnitude in the background region. Indicates how large the estimated force in the background is relative to the average magnitude of the actual traction force. DTMEB: Represents the deviation of traction force magnitude in the extracellular background region. Indicates how large the estimated force in the extracellular background is relative to the average magnitude of the actual traction force. DTMIB: Represents the deviation of traction force magnitude in the intracellular background region. Indicates how large the estimated force in the intracellular background is relative to the average magnitude of the actual traction force. For all metrics, a value closer to zero indicates better performance.

Incorporating morphological information introduces strong spatial constraints on where forces are allowed to occur, making it an effective method to guide biologically plausible predictions. However, this also imposes new requirements on dataset construction, as the distribution and localization of forces within the cell must be coherent with the given shape. Arbitrary force placement can hinder learning. To address this, we introduced biologically informed constraints that localize force generation to specific regions such as pseudopodia and focal adhesions, where traction forces are known to originate. While this strategy improves physical fidelity, it may not generalize across all cell types and physiological states, and caution should be exercised when extending the model to more complex or dynamic cellular behaviors.

Ultimately, the use of cell morphology in traction force inference is likely to become increasingly powerful as our understanding of the underlying biomechanics advances. Insights into the roles of actin filaments, adhesion complexes, cell polarity, and protrusion dynamics will enable more precise modeling of spatial constraints. Such developments are expected to further improve estimation accuracy and bridge the gap between mechanical modeling and biological function.

## 5 Conclusion

In this study, we proposed a machine learning framework for TFM that integrates cell morphology and biophysical mechanisms underlying force generation. By modeling the spatial origin, direction, and magnitude of traction forces based on cellular structures such as stress fibers and focal adhesions, we constructed biologically plausible training datasets. These datasets enabled a CNN to learn the inverse mapping from bead displacement to traction force with improved accuracy.

We demonstrated that incorporating cell shape information significantly enhances force estimation, especially under low bead-density conditions. Furthermore, we introduced a loss function that accounts for both the magnitude and direction of traction vectors, leading to improved directional accuracy and suppression of false predictions in non-adherent regions. Evaluation using multiple metrics confirmed that our model achieves robust and biologically consistent performance across various observation settings.

This approach provides a scalable framework for non-invasive biomechanical analysis using machine learning, and underscores the importance of combining observational data with mechanistic modeling. Future extensions may include adapting the model to time-varying forces, dynamic morphologies, and broader cell types, further bridging the gap between biophysical theory and experimental data.

## References

[1] M Tessier-Lavigne and CS Goodman. The molecular biology of axon guidance. Science, 274(5290):1123–1133, 1996. doi: 10.1126/science.274.5290.1123.

[2] Q Ma, D Jones, PR Borghesani, RA Segal, T Nagasawa, T Kishimoto, RT Bronson, and TA Springer. Impaired B-lymphopoiesis, myelopoiesis, and derailed cerebellar neuron migration in CXCR4- and SDF-1-deficient mice. Proc. Natl. Acad. Sci. U. S. A., 95(16):9448–9453, 1998. doi: 10.1073/pnas.95.16.9448.

[3] Barry J Dickson. Molecular mechanisms of axon guidance. Science, 298(5600):1959–1964, 2002. doi: 10.1126/science.1072165.

[4] Karen Lai Wing Sun, James P Correia, and Timothy E Kennedy. Netrins: versatile extracellular cues with diverse functions. Development, 138(11):2153–2169, 2011. doi: 10.1242/dev.044529.

[5] Andrew D Luster, Ronen Alon, and Ulrich H von Andrian. Immune cell migration in inflammation: present and future therapeutic targets. Nat. Immunol., 6(12):1182–1190, 2005. doi: 10.1038/ni1275.

[6] Filip K Swirski, Matthias Nahrendorf, Martin Etzrodt, Moritz Wildgruber, Virna Cortez-Retamozo, Peter Panizzi, Jose-Luiz Figueiredo, Rainer H Kohler, Aleksey Chudnovskiy, Peter Waterman, Elena Aikawa, Thorsten R Mempel, Peter Libby, Ralph Weissleder, and Mikael J Pittet. Identification of splenic reservoir monocytes and their deployment to inflammatory sites. Science, 325(5940):612–616, 2009. doi: 10.1126/science.1175202.

[7] Peter Friedl, Joseph Locker, Erik Sahai, and Jeffrey E Segall. Classifying collective cancer cell invasion. Nat. Cell Biol., 14(8):777–783, 2012. doi: 10.1038/ncb2548.

[8] NV Krakhmal, MV Zavyalova, EV Denisov, SV Vtorushin, and VM Perelmuter. Cancer invasion: Patterns and mechanisms. Acta Naturae, 7(2):17–28, 2015. doi: 10.32607/20758251-2015-7-2-17-28.

[9] Dennis E Discher, Paul Janmey, and Yu-Li Wang. Tissue cells feel and respond to the stiffness of their substrate. Science, 310(5751):1139–1143, 2005. doi: 10.1126/science.1116995.

[10] M Radmacher, M Fritz, CM Kacher, JP Cleveland, and PK Hansma. Measuring the viscoelastic properties of human platelets with the atomic force microscope. Biophys. J., 70(1):556–567, 1996. doi: 10.1016/s0006-3495(96)79602-9.

[11] Marcos Penedo, Keisuke Miyazawa, Naoko Okano, Hirotoshi Furusho, Takehiko Ichikawa, Mohammad Shahidul Alam, Kazuki Miyata, Chikashi Nakamura, and Takeshi Fukuma. Visualizing intracellular nanostructures of living cells by nanoendoscopy-AFM. Sci. Adv., 7(52):eabj4990, 2021. doi: 10.1126/sciadv.abj4990.

[12] John L Tan, Joe Tien, Dana M Pirone, Darren S Gray, Kiran Bhadriraju, and Christopher S Chen. Cells lying on a bed of microneedles: an approach to isolate mechanical force. Proc. Natl. Acad. Sci. U. S. A., 100(4):1484–1489, 2003. doi: 10.1073/pnas.0235407100.

[13] Ingmar Schoen, Wei Hu, Enrico Klotzsch, and Viola Vogel. Probing cellular traction forces by micropillar arrays: contribution of substrate warping to pillar deflection. Nano Lett., 10(5):1823–1830, 2010. doi: 10.1021/nl100533c.

[14] M Dembo, T Oliver, A Ishihara, and K Jacobson. Imaging the traction stresses exerted by locomoting cells with the elastic substratum method. Biophys. J., 70(4):2008–2022, 1996. doi: 10.1016/s0006-3495(96)79767-9.

[15] Clarence E Chan and David J Odde. Traction dynamics of filopodia on compliant substrates. Science, 322(5908):1687–1691, 2008. doi: 10.1126/science.1163595.

[16] Margaret L Gardel, Benedikt Sabass, Lin Ji, Gaudenz Danuser, Ulrich S Schwarz, and Clare M Waterman. Traction stress in focal adhesions correlates biphasically with actin retrograde flow speed. J. Cell Biol., 183(6):999–1005, 2008. doi: 10.1083/jcb.200810060.

[17] Jonathan Stricker, Benedikt Sabass, Ulrich S Schwarz, and Margaret L Gardel. Optimization of traction force microscopy for micron-sized focal adhesions. J. Phys. Condens. Matter, 22(19):194104, 2010. doi: 10.1088/0953-8984/22/19/194104.

[18] Michinori Toriyama, Satoshi Kozawa, Yuichi Sakumura, and Naoyuki Inagaki. Conversion of a signal into forces for axon outgrowth through Pak1-mediated shootin1 phosphorylation. Curr. Biol., 23(6):529–534, 2013. doi: 10.1016/j.cub.2013.02.017.

[19] Claude N Holenstein, Unai Silvan, and Jess G Snedeker. High-resolution traction force microscopy on small focal adhesions improved accuracy through optimal marker distribution and optical flow tracking. Sci. Rep., 7(1):41633, 2017. doi: 10.1038/srep41633.

[20] I Jasiuk, PY Sheng, and E Tsuchida. A spherical inclusion in an elastic half-space under shear. J. Appl. Mech., 64(3):471–479, 1997. doi: 10.1115/1.2788917.

[21] Volodymyr I Kushch. Elastic ellipsoidal inhomogeneity with imperfect interface: Complete displacement solution in terms of ellipsoidal harmonics. Int. J. Solids Struct., 166:83–95, 2019. doi: 10.1016/j.ijsolstr.2019.02.007.

[22] Yunfei Huang, Christoph Schell, Tobias B Huber, Ahmet Nihat Ş imşek, Nils Hersch, Rudolf Merkel, Gerhard Gompper, and Benedikt Sabass. Traction force microscopy with optimized regularization and automated bayesian parameter selection for comparing cells. Sci. Rep., 9(1):539, 2019. doi: 10.1038/s41598-018-36896-x.

[23] Ryosuke Fujikawa, Chika Okimura, Satoshi Kozawa, Kazushi Ikeda, Naoyuki Inagaki, Yoshiaki Iwadate, and Yuichi Sakumura. Bayesian traction force estimation using cell boundary-dependent force priors. Biophys. J., 122(23):4542–4554, 2023. doi: 10.1016/j.bpj.2023.10.032.

[24] Yu-Li Wang and Yun-Chu Lin. Traction force microscopy by deep learning. Biophys. J., 120(15):3079–3090, 2021. doi: 10.1016/j.bpj.2021.06.011.

[25] Honghan Li, Daiki Matsunaga, Tsubasa S Matsui, Hiroki Aosaki, Genki Kinoshita, Koki Inoue, Amin Doostmohammadi, and Shinji Deguchi. Wrinkle force microscopy: a machine learning based approach to predict cell mechanics from images. Commun. Biol., 5(1):361, 2022. doi: 10.1038/s42003-022-03288-x.

[26] Felix S Kratz, Lars Möllerherm, and Jan Kierfeld. Enhancing robustness, precision, and speed of traction force microscopy with machine learning. Biophys. J., 122(17):3489–3505, 2023. doi: 10.1016/j.bpj.2023.07.025.

[27] Thomas D Pollard and Gary G Borisy. Cellular motility driven by assembly and disassembly of actin filaments. Cell, 112(4):453–465, 2003. doi: 10.1016/s0092-8674(03)00120-x.

[28] A Ponti, M Machacek, SL Gupton, CM Waterman-Storer, and G Danuser. Two distinct actin networks drive the protrusion of migrating cells. Science, 305(5691):1782– 1786, 2004. doi: 10.1126/science.1100533.

[29] Olaf Ronneberger, Philipp Fischer, and Thomas Brox. Unet: Convolutional networks for biomedical image segmentation. In Medical Image Computing and Computer-Assisted Intervention (MICCAI), pages 234–241. Springer, 2015. doi: 10.1007/978-3-319-24574-4_28.

[30] W Helfrich. Elastic properties of lipid bilayers: theory and possible experiments. Z. Naturforsch. C, 28(11):693–703, 1973. doi: 10.1515/znc-1973-11-1209.

[31] W Helfrich. The size of bilayer vesicles generated by sonication. Physics Letters A, 50:115–116, 1974. doi: 10.1016/0375-9601(74)90899-8.

[32] W Helfrich. Blocked lipid exchange in bilayers and its possible influence on the shape of vesicles. Z. Naturforsch. C, 29C(9-10):510–515, 1974. doi: 10.1515/znc-1974-9-1010.

[33] Angelo Angonezi, Fernanda Oliveira, Juliano Faccioni, Camila Cassel, Débora Santos de Sousa, Samlai Vedovatto, Guido Lenz, Cláudio Jung, and Lucas Kirsten. Oriented cell dataset (ocd). IEEE Dataport, January 2024. doi:10.21227/3rk2-b882.

[34] Carsen Stringer, Tim Wang, Michalis Michaelos, and Marius Pachitariu. Cellpose: a generalist algorithm for cellular segmentation. Nat. Methods, 18(1):100–106, 2021. doi: 10.1038/s41592-020-01018-x.

[35] Alec Radford, Luke Metz, and Soumith Chintala. Unsupervised Representation Learning with Deep Convolutional Generative Adversarial Networks. arXiv preprint arXiv:1511.06434, 2015. doi: 10.48550/arXiv.1511.06434.

[36] Anne J Ridley, Martin A Schwartz, Keith Burridge, Richard A Firtel, Mark H Ginsberg, Gary Borisy, J Thomas Parsons, and Alan Rick Horwitz. Cell migration: integrating signals from front to back. Science, 302(5651):1704–1709, 2003. doi: 10.1126/science.1092053.

[37] Ronen Zaidel-Bar, Christoph Ballestrem, Zvi Kam, and Benjamin Geiger. Early molecular events in the assembly of matrix adhesions at the leading edge of migrating cells. J. Cell Sci., 116(Pt 22):4605–4613, 2003. doi: 10.1242/jcs.00792.

[38] Sari Tojkander, Gergana Gateva, and Pekka Lappalainen. Actin stress fibers–assembly, dynamics and biological roles. J. Cell Sci., 125(Pt 8):1855–1864, 2012. doi: 10.1242/jcs.098087.

[39] L.D. Landau and E.M. Lifshitz. Theory of Elasticity. Second Revised and Enlarged. Pergamon Press Ltd, 1970.

[40] Hao Wang, Yitong Wang, Zheng Zhou, Xing Ji, Dihong Gong, Jingchao Zhou, Zhifeng Li, and Wei Liu. CosFace: Large margin cosine loss for deep face recognition. In 2018 IEEE/CVF Conference on Computer Vision and Pattern Recognition. IEEE, 2018. doi: 10.1109/cvpr.2018.00552.

